# Long-read sequencing and profiling of RNA-binding proteins reveals the pathogenic mechanism of aberrant splicing of an *SCN1A* poison exon in epilepsy

**DOI:** 10.1101/2023.05.04.538282

**Authors:** Hannah C Happ, Patricia N Schneider, Jung Hwa Hong, Eleanor Goes, Masha Bandouil, Carina G. Biar, Aishwarya Ramamurthy, Fairlie Reese, Krysta Engel, Sarah Weckhuysen, Ingrid E Scheffer, Heather C Mefford, Jeffrey D Calhoun, Gemma L Carvill

## Abstract

Pathogenic loss-of-function *SCN1A* variants cause a spectrum of seizure disorders. We previously identified variants in individuals with *SCN1A*-related epilepsy that fall in or near a poison exon (PE) in *SCN1A* intron 20 (20N). We hypothesized these variants lead to increased PE inclusion, which introduces a premature stop codon, and, therefore, reduced abundance of the full-length *SCN1A* transcript and Na_v_1.1 protein. We used a splicing reporter assay to interrogate PE inclusion in HEK293T cells. In addition, we used patient-specific induced pluripotent stem cells (iPSCs) differentiated into neurons to quantify 20N inclusion by long and short-read sequencing and Na_v_1.1 abundance by western blot. We performed RNA-antisense purification with mass spectrometry to identify RNA-binding proteins (RBPs) that could account for the aberrant PE splicing. We demonstrate that variants in/near 20N lead to increased 20N inclusion by long-read sequencing or splicing reporter assay and decreased Na_v_1.1 abundance. We also identified 28 RBPs that differentially interact with variant constructs compared to wild-type, including SRSF1 and HNRNPL. We propose a model whereby 20N variants disrupt RBP binding to splicing enhancers (SRSF1) and suppressors (HNRNPL), to favor PE inclusion. Overall, we demonstrate that *SCN1A* 20N variants cause haploinsufficiency and *SCN1A*-related epilepsies. This work provides insights into the complex control of RBP-mediated PE alternative splicing, with broader implications for PE discovery and identification of pathogenic PE variants in other genetic conditions.

## Introduction

*SCN1A* (MIM: 182389) encodes the neuronal voltage-gated sodium channel subunit Na_v_1.1. Pathogenic loss-of-function (LOF) variants in *SCN1A* cause a spectrum of seizure disorders, ranging from febrile seizures and genetic epilepsy with febrile seizures plus (GEFS+) to severe developmental and epileptic encephalopathies (DEEs), including Dravet syndrome. Individuals with Dravet syndrome typically present with febrile seizures around six months of age, develop multiple seizure types and drug-resistant epilepsy, and are at high risk of sudden death in epilepsy. Approximately 90% of individuals with Dravet syndrome have *de novo SCN1A* truncation or LOF missense variants^1,2^.

A prior sequencing study of highly conserved *SCN1A* intronic regions in a cohort of 640 individuals with an unsolved DEE identified five variants in five individuals with *SCN1A*-related phenotypes^3^. These variants fall in or near a poison exon (PE) in *SCN1A* intron 20. PEs are small, highly conserved exons that lead to the introduction of a premature truncation codon (PTC). When a PE is incorporated into a mature mRNA transcript by alternative splicing, this can trigger mRNA degradation via nonsense-mediated decay (NMD). Since this transcript does not yield a functional protein, we refer to this as unproductive splicing^4,5^. Although the role of PEs is not fully understood, they likely function as a means of tightly and rapidly regulating gene expression within a cell. For example, genes encoding RNA-binding proteins (RBPs) are enriched for PEs, and the unproductive splicing that results from PE inclusion is used to regulate gene expression^6,7^. Moreover, alternative splicing of PEs can be cell type- or tissue-specific and is widely used to modulate gene expression during neurodevelopment^8,9^. This context-specific inclusion/exclusion of PEs is controlled by RBPs that regulate splicing by recognition of specific RNA sequence motifs. For example, RBFOX proteins regulate neuron-specific exclusion of a PE in *SCN8A*^*10*^. In addition, many alternatively spliced exons in mouse neurons are bound by RBPs PTBP1 and/or RBFOX1/2/3^9^, though the specific RBPs that control PE alternative splicing are not known. To date, the only disease-associated PE-specific splicing study demonstrated that disruption of a PTBP1 binding site in *FLNA*, which leads to increased inclusion of a PE, can cause periventricular nodular heterotopia in humans^9^.

We hypothesized that pathogenic variants in or near a PE in *SCN1A* intron 20, known as 20N, lead to increased inclusion of 20N and cause *SCN1A-*related phenotypes. We previously described five individuals with variants in or near 20N (**Supplementary Table S1**); four individuals had Dravet syndrome, and one had Febrile Seizures Plus (FS+)^3^ (**Figure 1**). We proposed that these variants lead to increased 20N inclusion in the mature transcript, resulting in degradation of the *SCN1A* transcript and reduced abundance of Na_v_1.1. This degradation of the *SCN1A* 20N-containing transcript is facilitated by NMD, as shown in human neural precursor cells (ReNcells), where inhibiting protein translation (and thus, NMD) with cycloheximide leads to increased abundance of the PE containing transcript^5,11^. A preliminary functional study using a splicing reporter assay in both A549 and K562 cells showed that three of the five variants in or near 20N lead to increased 20N inclusion^3^. However, this study was limited to the splicing assay, functional experimentation was not performed in patient-specific cells, and SCN1A haploinsufficiency was not demonstrated. Moreover, the RBPs that regulate putative aberrant PE splicing are unknown. Therefore, in this study, we investigate 20N inclusion in patient-specific induced pluripotent stem cell (iPSC)-derived neuronal models and interrogate the mechanism by which the genetic variants promote 20N inclusion using a splicing reporter in HEK293T cells. Our work informs the pathogenic mechanism underlying aberrant PE splicing and aids future RNA therapeutics design as well as understanding how PEs may play a broader role in disease.

**Figure 1.**
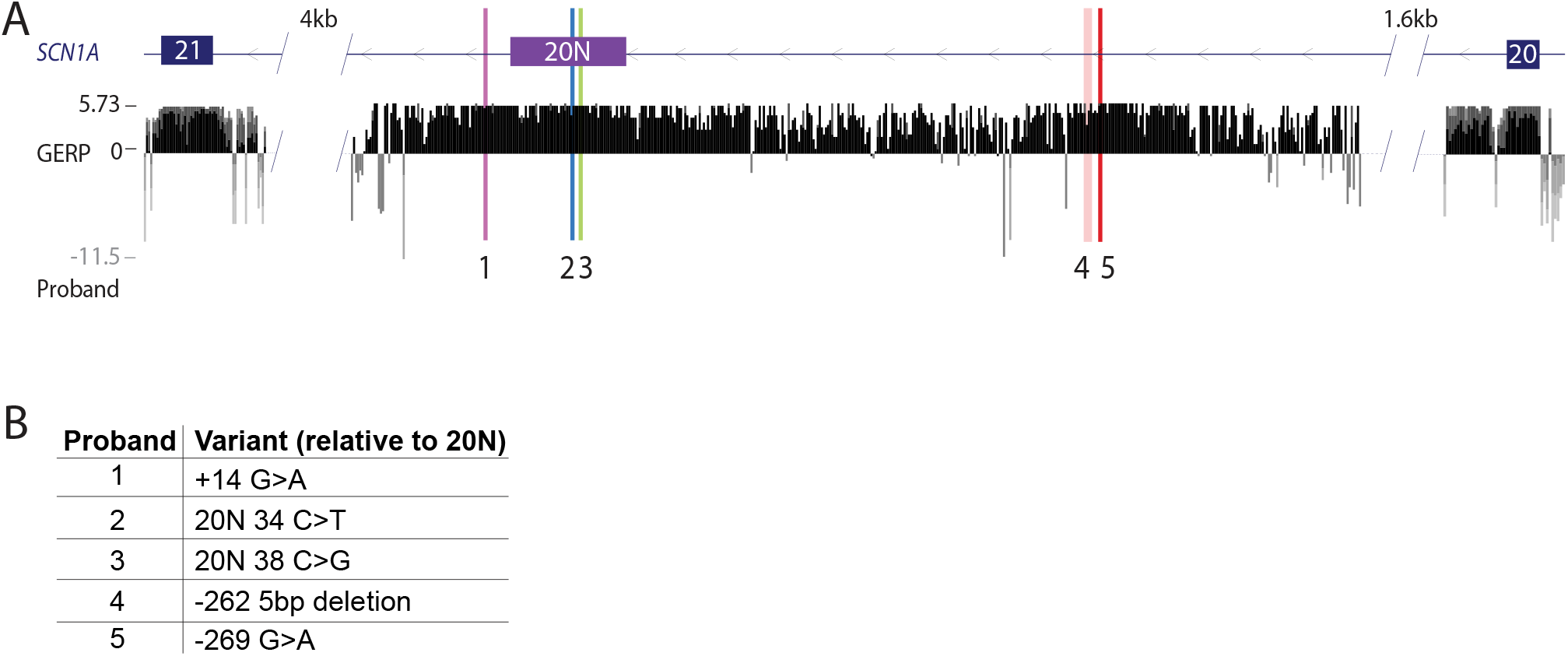
Pathogenic variants in and around *SCN1A* poison exon (PE) 20N in 5 individuals with *SCN1A-*associated epilepsy. **(A)** *SCN1A* PE 20N is located between canonical exons 20 and 21. The orientation of *SCN1A* is shown from right to left because it codes off the negative strand of DNA. 20N and the 5 variants are located in a highly conserved region of intron 20, indicated by positive Genomic Evolutionary Rate Profiling (GERP) scores. **(B)** Of the five previously identified variants^3^, two are located within 20N, variant 1 is located 14 bp downstream, variants 2 and 3 are located in 20N, and variants 4 and 5 are located 262 and 269 bp upstream of 20N, respectively. Variants 1, 2, 3, and 5 are single nucleotide variants, and variant 4 is a five bp deletion.

## Results

### Specific variants in or near SCN1A 20N PE affect 20N inclusion in a splicing reporter assay

To test the effect of the five patient-specific variants on 20N splicing, we transfected HEK293T cells with the 20N splicing reporter construct and used RT-qPCR to show variants 2 and 3 lead to a clear increase in 20N inclusion compared to wild-type (WT) for both the 20N:Exon21 and Exon20:20N assay (**Supplementary Table S2 and Figure S1**). Conversely, for the variant 1 construct, only the 20N:Exon21 assay showed increased 20N inclusion. This discrepancy could be the result of mutually exclusive inclusion of 20N and exon 20 or a technical limitation of the splicing reporter system. Variants 4 and 5 did not lead to increased inclusion of 20N compared to WT in this splicing reporter assay.

### Targeted long-read sequencing reveals increased inclusion of SCN1A 20N in patient-specific iPSC-derived neurons

To interrogate the discrepancy in 20N inclusion observed for variant 1 and the lack of increased 20N inclusion shown for variant 4 and 5 constructs, we utilized targeted long-read *SCN1A* RNA sequencing in patient-specific neurons. Long-read RNA sequencing captures entire transcripts in a single read, allowing for robust detection of isoform-specific expression. Since *SCN1A* is not expressed in fibroblasts or induced pluripotent stem cells (iPSCs), we differentiated patient-specific iPSCs into neurons. Sequencing of the entire transcript enabled us to confidently conclude that 20N is a cassette exon (spliced to exons 20 and 21) in the majority of transcripts (**Figure 2A**). To account for variability in neuron maturation, we normalized 20N inclusion to *SCN3A* and *SCN8A* expression from short-read whole transcriptome sequencing of the same samples. These two transcripts were selected as markers of maturation as the reciprocal relationships of these two subunits is well known, with *SCN3A* being expressed very early in development, and *SCN8A* later^12^. We demonstrate a significant increase in 20N inclusion in neurons from probands 1, 4, and 5 compared to control neurons (P-adjusted = 0.0003, 0.0054, 0.0045, respectively) (**Figure 2B**). Overall, we demonstrate a twofold increase in 20N inclusion in proband vs control neurons (P = 0.0004) (**Figure 2C**). We did not have access to iPSCs from probands 2 and 3 to perform this transcript analysis for their variants. Collectively, our results indicate that in addition to variants 2 and 3, variants 1, 4, and 5 also lead to increased inclusion of 20N in the mature *SCN1A* transcript in a highly relevant cell type.

**Figure 2.**
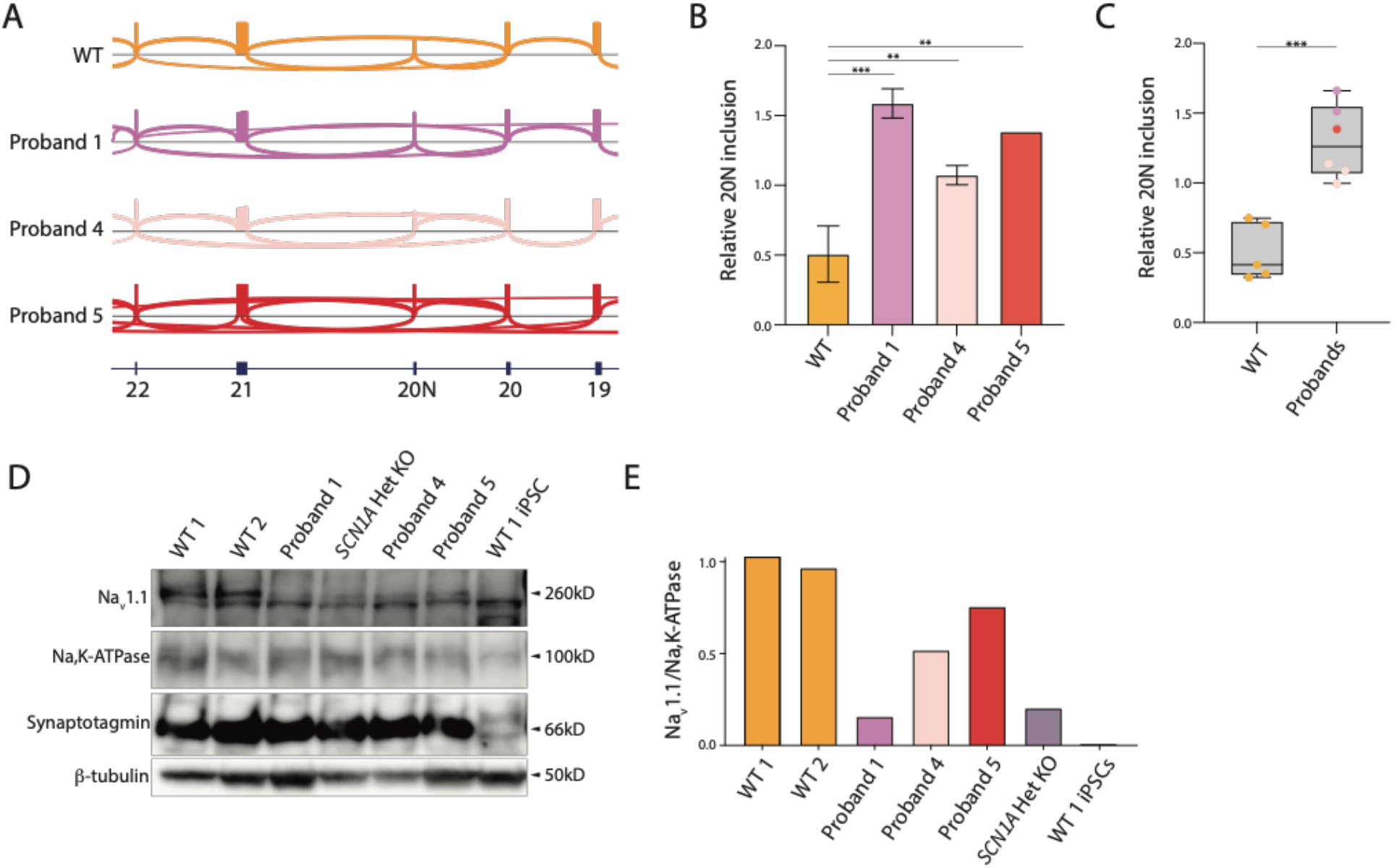
Pathogenic variants in and around *SCN1A* poison exon 20N lead to increased 20N inclusion and reduced abundance of Na_v_1.1 in proband-specific iPSC-derived neurons. Proband and control iPSCs were differentiated into neurons to perform long-read sequencing and protein analysis. **(A)** Sashimi plots illustrate alternative splicing events identified by long-read RNA sequencing. Vertical bars indicate exons, and the height of the bars is proportionate to the read depth supporting the inclusion of an exon. Horizontal arcs illustrate splicing junctions that are supported by the sequencing reads. For visualization purposes, we highlight the region of *SCN1A* between exons 19 and 22, although individual reads capture the entire transcript. Sashimi plots are representative plots from one biological replicate. **(B)** We quantified the relative abundance of transcripts that include 20N for each sample. Abundance was calculated as a percentage of inclusion transcripts, normalized for read depth and cell maturation. Cell maturation was determined by the relative normalized expression of *SCN3A* and *SCN8A*. Two WT cell lines with the same genotype were combined for statistical analysis. Data are presented as the mean ± SD. For comparisons, an ordinary one-way ANOVA with post-hoc test was performed on WT-variant pairs (*P-adjusted ≤ 0.05; **P ≤ 0.01; ***P<0.001). Biologic replicates *n* = 1-3. **(C)** Relative abundance of 20N inclusion transcripts grouped by genotype. For comparisons, student’s unpaired two-tailed t-test was performed (***P<0.001). **(D)** Proband iPSC-derived neurons show less Na_v_1.1 protein as detected by Western blot. Samples were prepared by membrane protein isolation, so Na,K-ATPase is used as a loading control. The sample in lane 4 serves as a positive control for haploinsufficiency as this lysate was prepared from neurons derived from an individual with Dravet syndrome who carries a heterozygous upstream deletion of the *SCN1A* transcription start site. Lysate from WT iPSCs is included as a negative control as iPSCs do not express *SCN1A*. Biologic replicates *n* = 1. **(E)** ImageJ quantification of Western blot bands shows reduced abundance of Na_v_1.1 in neurons from probands 1, 4, and 5 as well as the haploinsufficiency control cells compared to WT.

### Increased inclusion of SCN1A 20N in neurons causes Na_v_1.1 haploinsufficiency

The inclusion of 20N in a transcript introduces a PTC (**Supplementary Figure S2**), which can target the transcript for NMD. We therefore hypothesized that the increased unproductive splicing in patients with variants 1-5 was sufficient to cause Na_v_1.1 haploinsufficiency in neurons. Western blot analysis of iPSC-derived neurons from probands 1, 4, and 5 showed an 84%, 48%, and 24% decreased abundance of Na_v_1.1 compared to WT neurons (**Figure 2D-E**). Cells from a haploinsufficiency control, carrying an upstream heterozygous deletion of the *SCN1A* transcription start site^3^, showed an 80% reduction in Na_v_1.1 abundance (**Figure 2D-E**). Ultimately, these results show that variants 1, 4, and 5 lead to a reduction in Na_v_1.1 protein, which is consistent with the phenotypes of Dravet syndrome and FS+ in the individuals carrying these variants.

### 20N variants alter RBP binding

Alternative splicing is controlled by RBPs that bind to specific sequence motifs in mRNA. We proposed that patient variants cause aberrant 20N splicing by altering RBP binding. Therefore, we utilized RNA antisense purification with mass spectrometry (RAP-MS) to identify RBPs that directly interact with the 20N region (**Figure 3A)**. We performed RAP-MS using WT, variant 1, and variant 2 constructs to identify differentially bound RBPs as we reasoned that variants 2 and 3, which are four base pairs apart, likely affect the same RBP-binding motifs. Overall, we identified 169 proteins bound to at least one sample (**Supplementary File 1**). 64 proteins were excluded from further analysis as they were identified in either the GFP-plasmid or untransfected controls and were thus likely non-specific. We prioritized 40 proteins with known RNA-binding activity. Of these RBPs, 12 are present in all three samples; 28 are differentially bound to WT and at least one variant construct (**Figure 3B**).

**Figure 3.**
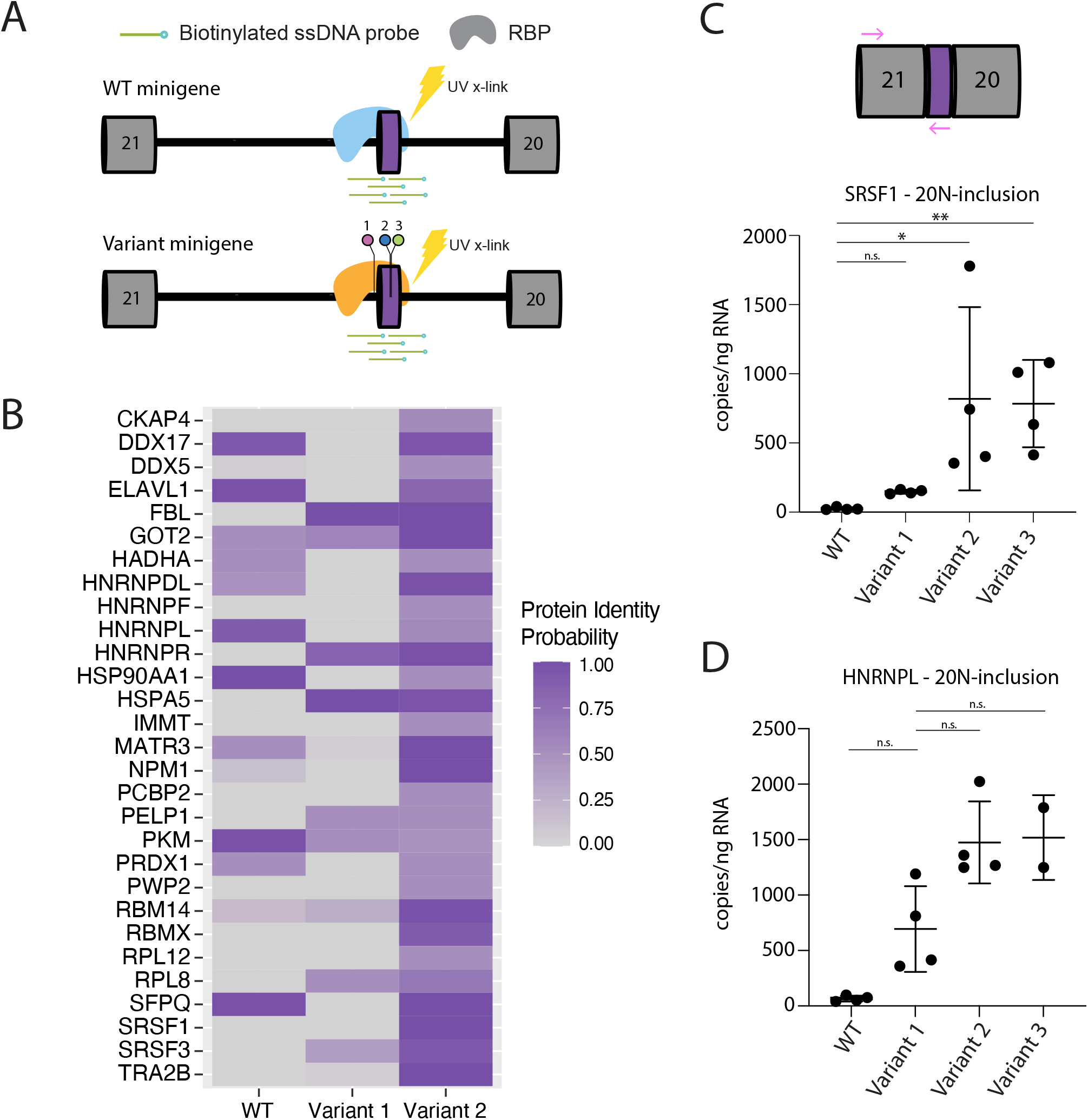
Identification of RNA-binding proteins (RBPs) that bind to WT and variant *SCN1A* splicing reporter constructs. **(A)** Splicing reporter constructs containing either WT sequence, variant 1 or variant 2 were transfected into HEK293T cells. RNA-RBP interactions were covalently captured by UV cross-linking constructs were hybridized to biotinylated DNA probes and captured with streptavidin beads. Variants 2 and 3 are four bp apart and likely affect the same RBP-binding motifs, therefore, we performed RAP-MS using WT, variant 1, and variant 2 constructs. **(B)** Proteomics analysis was performed by mass spectrometry to identify RBPs that bind the reporter constructs. Detected peptide sequences are matched against a protein database to identify the protein of origin, which is represented by a Protein Identity Probability score (scale 0-1). Lower scores indicate a protein is absent in the sample; higher scores indicate a protein is present. 28 RBPs are differentially bound to variant splicing constructs compared to WT. Scores represent the average of two biologic replicates. **(C and D)** RIP was performed with the same splicing reporter constructs used for RAP-MS with the addition of a variant 3 construct. RNA-RBP interactions were captured by UV cross-linking, and immunoprecipitations were carried out with antibodies for SRSF1 (C) or HNRNPL (D). Following RNA extraction and reverse transcription, droplet-digital PCR with 20ng RNA was used to quantify 20N-inclusion transcripts bound to SRSF1 (C) or HNRNPL (D). Kruskal-Wallis comparisons with Dunn’s post-hoc test were performed on WT-variant pairs (SRSF1) or variant 1-WT/variant pairs (HNRNPL) (*P-adjusted ≤ 0.05; **P ≤ 0.01). Biological replicates *n* = 4 or *n* =2 (HNRNPL variant 3), technical replicates *n* = 2. Data are presented as the mean ± SD.

We selected SRSF1 and HNRNPL for validation via RNA immunoprecipitation (RIP) based on their established roles as regulators of alternative splicing, where SRSF1 tends to promote, and HNRNPL tends to suppress alternative splicing^13,14^. Of the RNA bound to SRSF1, there was a statistically significant increase in the SRSF1-bound 20N-inclusion transcript for variant 2 and 3 compared to WT and variant 1 (**Figure 3C**). Conversely, the HNRNPL-bound 20N-inclusion transcript is less abundant in the variant 1 sample (**Figure 3D)** compared to variants 2 and 3. The SRSF1/HNRNPL-bound 20N-inlcusion WT transcript was much lower than all variants, possibly due to the lower inclusion of 20N overall (**Supplementary Figure S1**). Moreover, the canonical transcript in general had much lower abundance, possibly due to abundance and PCR efficiency (**Supplementary Figure S4**). Together, these findings validate the RAP-MS findings whereby PE variants alter RBP binding to 20N.

## Discussion

Haploinsufficiency of *SCN1A* is associated with a well-established epilepsy phenotypic spectrum, ranging from Dravet syndrome to GEFS+. These epilepsies are typically caused by pathogenic loss-of-function missense or truncation variants in the coding region of *SCN1A*. We previously identified five likely pathogenic variants in (*n*=2) or near (*n*=3) a PE in *SCN1A* intron 20 in individuals with Dravet syndrome or FS+^3^. Here we show that all five variants lead to increased inclusion of 20N in *SCN1A* and, where patient-derived cells were available, these variants lead to reduced abundance of Na_v_1.1 protein. We identify RBPs that differentially interact with variant transcripts and drive aberrant PE inclusion and develop an early conceptual model for the *SCN1A* PE alternative splicing that may be used as a framework for future work.

The five probands in this study have phenotypes consistent with *SCN1A* haploinsufficiency; however, additional functional analysis was needed to demonstrate Na_v_1.1 loss and determine the underlying mechanism. Collectively, using two complementary techniques, we show that all five variants lead to increased 20N inclusion. Variants 1, 2, and 3 led to increased inclusion of 20N in HEK293T cells, mirroring previous findings in K562 and A549 cells. In contrast to the clear conclusions from variants 2 and 3, variant 1 showed increased inclusion using only one primer set and variants 4 and 5 showed no increase in 20N inclusion. However, we demonstrate that 20N is included in significantly more transcripts in neurons from probands 1, 4, and 5 compared to WT iPSC-derived neurons; furthermore we show a reduction in Na_v_1.1 abundance, consistent with *SCN1A* haploinsufficiency. The increased 20N inclusion and introduction of a PTC likely leads to degradation of 20N-inclusion transcripts, yielding the concomitant reduction in protein. Previous studies in mice and human ReNcells have demonstrated that 20N inclusion leads to NMD, and inhibition of NMD with cycloheximide results in even greater abundance of 20N inclusion transcripts^5,11^. Our work also highlights the benefit of using patient-specific iPSC-derived cells, particularly to study the transcription landscape of genes that are highly cell-type specific. This model is especially powerful for neurodevelopmentally-regulated genes because it allows otherwise inaccessible cell types to be studied. Ultimately, utilizing these cells was critical to determining the pathogenic mechanism of haploinsufficiency due to increased inclusion of 20N especially for variants 4 and 5 which failed to increased 20N inclusion in the splice reporter assay in HEK293T, A549, and K562 cell lines.

Determining the RBPs that control splicing of PEs in general, and *SCN1A* specifically, is critical to our understanding of how PEs are alternatively spliced and how their splicing is disrupted in the presence of variants. Here we identified 28 RBPs that differentially interact with splicing reporters that contain either the WT sequence or variants 1 or 2. Overall, these results highlight that alternative splicing is highly complex and controlled by many RBPs and there are still many future studies needed to unravel this complexity. We focused here initially on those RBPs that disrupted known consensus motifs and play well-established roles in alternative splicing of cassette exons. For example, in one scenario RBPs can bind specifically to either intronic or exonic regions to act as alternative splicing silencers or enhancers (**Figure 4**). Two well-studied antagonistic RBP enhancers/silencers are the serine/arginine-rich (SR) proteins such as SRSF1 which tend to bind enhancers to promote the inclusion of an exon, while heterogeneous nuclear ribonuclear (HNRNP) proteins tend to bind suppressors to prevent exon inclusion (**Figure 4A**). Our work demonstrates that variant 1 weakens the binding of HNRNPL, likely by disrupting a CA-rich motif, decreasing affinity for a putative intronic splicing suppressor and promoting 20N inclusion (**Figure 4B and Supplementary Figure S5A**). Meanwhile, variants 2 and 3 enhance the binding of SRSF1 to a putative splicing enhancer, promoting 20N inclusion (**Figure 4C and Supplementary Figure S5B**). SRSF1 tends to bind GA-rich motifs, which makes the interpretation of variants 2 and 3 challenging, because both variants replace a G at the mRNA level. Interestingly, SRSF1 RNA recognition motif 1 can preferentially bind CN motifs, and disruption of a CA motif in SMN1 exon7 activates alternative splicing of an exon in SMN1^15^. This may explain how variant 3, which causes a G-to-C (mRNA) change, enhances SRSF1 binding. As we have demonstrated that SRSF1 binding is low in both WT and variant 1 context, which maintain G nucleotides at both positions, this highlights that the current understanding of RBP motifs is limited and will improve over time. Ultimately, variant disruption of 20N splicing is likely more complex than can be encompassed by this working model, particularly because RBPs, including SRSF1 and HNRNPL can have numerous functions and combinatorial consequences that are context-dependent^16^. However, it provides an important framework for considering the pathogenic mechanism of PE variants, particularly variants 1, 2, and 3, and informs how we might further interrogate the functional consequences of variants that affect PEs and other alternative splicing events in the future.

**Figure 4.**
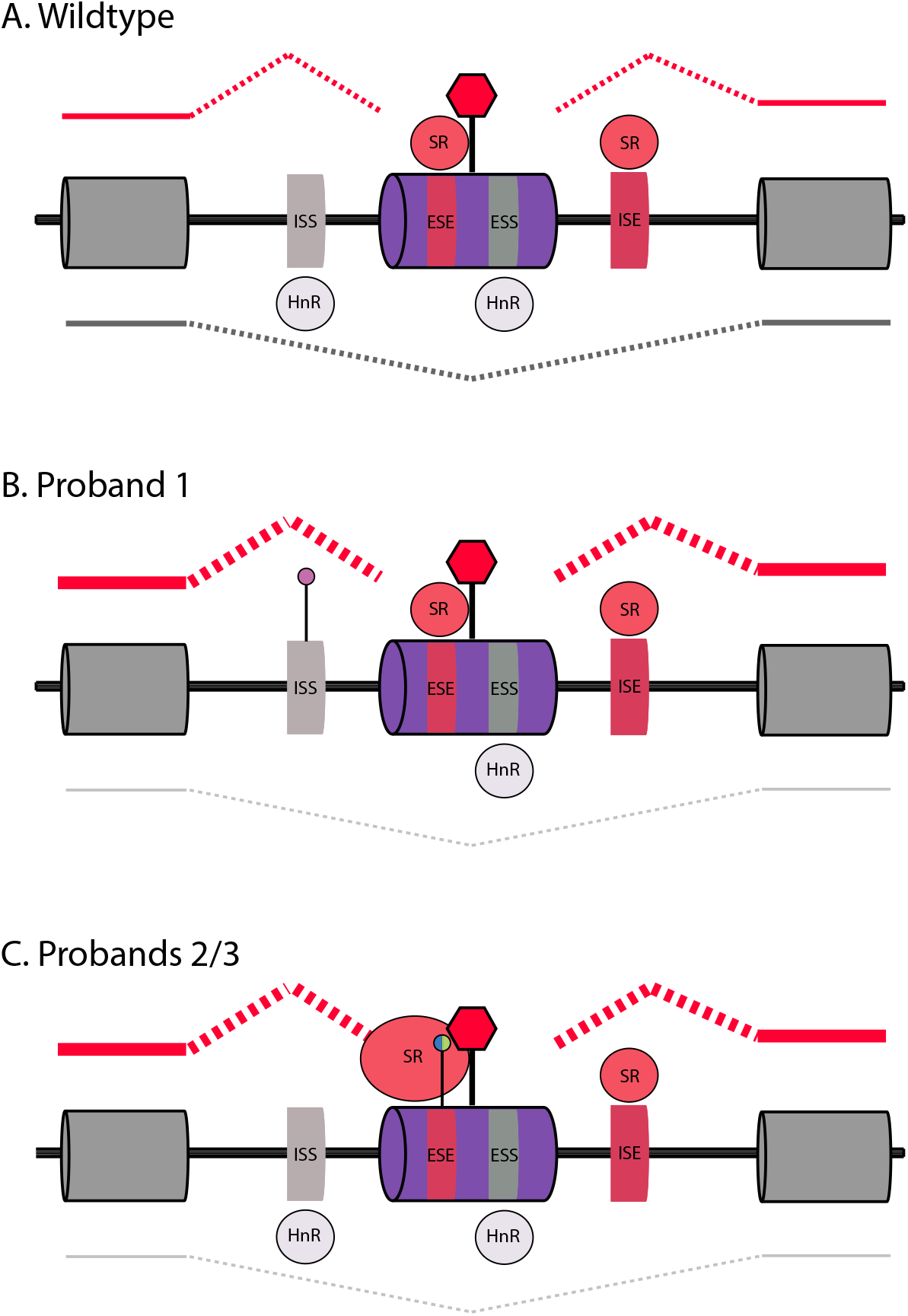
An early conceptual model of how variants disrupt RNA binding protein (RBP) binding to intronic and exonic splice enhancers/suppressors of the *SCN1A* poison exon. **(A)** SR RBPs can bind intronic/exonic splicing enhancers and promote alternate exon inclusion. HNRNP RBPs can bind intronic/exonic suppressor elements and prevent alternative exon splicing. **(B)** Variant 1 may prevent binding of HNRNPL to a putative ISS, weakening the suppressor element and therefore favoring PE inclusion. **(C)** Variants 2 and 3 enhance SRSF1 binding to a putative exonic ESE, strengthening the enhancer element and associated PE inclusion. ISS: intronic splicing suppressor; ISE: intronic splicing enhancer; ESS: exonic splicing suppressor; ESE: exonic splicing enhancer; SR: serine/arginine-rich protein; HnR: heterogeneous nuclear ribonucleoprotein.

We have highlighted the utility of splicing reporters to both assess how variants can disrupt splicing, especially when patient-specific cells are not available, and assay RBP control of splicing at a scale that supports proteomics analysis. However, this approach has limitations, including that variants 4 and 5 clearly alter 20N splicing in neurons but not in the splicing reporter assay. RBP control of alternative splicing varies by cell/tissue type and developmental stage. Variants 4 and 5 likely alter 20N splicing in neurons by altering the binding of RBPs present in neurons but absent in A549, K562, and HEK293T cells. For example, variants 4 and 5 create AU-rich motifs (**Supplementary Figure S5C**), which are recognized by neuronal Elav-like (Hu) proteins ELAVL3 (HuC) and ELAVL4 (HuD)^17^, which are expressed in neurons but not in the three cell types we investigated here or previously^3^. Furthermore, many RBPs are sensitive to the three-dimensional structure RNA context^18^, which cannot be fully recapitulated by the splicing reporter construct.

Aberrant PE splicing is not limited to *SCN1A* and indeed may be an important mechanism that underpins some neurodevelopmental disorders (NDDs) with unknown etiology. For example, variants near *SCN2A* and *SCN8A* PEs have also been described in individuals with epilepsy^19^. Furthermore, pathogenic variants that affect PEs have been implicated in diseases such as periventricular nodular heterotopia^9^, craniofacial microsomia^20^, and a rare recessive neurodegenerative disorder^21^. However, identifying these variants is difficult for several reasons, including the fact that it is difficult to identify the PEs themselves. The inclusion of PEs can be cell-type specific and/or developmentally regulated, and their inclusion naturally leads to the degradation of the transcript via NMD, thus detecting PEs by short-read RNA-sequencing is challenging. Future studies utilizing long-read sequencing to capture PE-inclusion transcripts in multiple cell types and across development will be an important tool. As we show, PE-inclusion transcripts can be identified without inhibiting NMD, but it is also likely that some PEs are included at much lower frequencies and could be missed. This problem can be addressed by sequencing very deeply or by utilizing techniques to inhibit NMD. Of course, identifying PEs is only part of the equation; we also need to be able interpret candidate variants, particularly as we show that disease-causing variants can reside within the PE but also hundreds of base pairs away. Here we would also benefit from tools that integrate empirical evidence of how RBPs control context-specific alternative splicing events with machine learning algorithms to predict PE splicing and, in turn, how variants may disrupt these splicing events.

Overall, we confirm that variants in and around an *SCN1A* PE cause increased 20N inclusion and the subsequent transcript degradation leads to haploinsufficiency, which underlie the phenotypes of the individuals carrying these variants. This work begins to unravel the complex control of RBPs on alternative splicing of PEs, which has broader implications for PE discovery and identification of pathogenic PE variants implicated in other genetic conditions, as well as translational therapeutic implications. Given their role in transcriptional regulation, PEs are potential targets for antisense oligonucleotide (ASO) therapies. ASOs can bind to a transcript and modulate splicing of a PE into the mature transcript. Given that *SCN1A* 20N is included at low levels in WT cells, targeting naturally occurring 20N inclusion with ASOs can yield increased splicing of the canonical transcript and a subsequent increase in protein abundance. This has been shown in pre-clinical studies using a mouse model of Dravet syndrome^11^ and is in clinical trials as a therapy for individuals with *SCN1A* haploinsufficiency caused by LOF missense or truncating variants. Understanding the mechanisms by which PE splicing is controlled will also aid in the identification of PEs, providing potential new therapeutic targets for genetic disorders. Identifying the RBPs that control PE splicing is useful for both surveying their binding sites for unidentified PEs and the potential to target genes that cause disease by gain-of-function, as ASOs could be used to promote binding of RBPs that lead to PE inclusion and decrease of gain-of-function (GOF) protein products. Ultimately, this study lays the groundwork for developing a framework to identify PEs via RBP binding and determine which variants in individuals with genetic conditions may disrupt PE splicing.

## Data availability

The datasets used and/or analyzed during the current study are included in this published article [and its supplementary information files] or are available from the corresponding author on reasonable request.

## Methods

### Study participants

All five probands were recruited as described previously^3^.

### SCN1A splicing reporter plasmids

pDESTsplice (Addgene #32484) splicing reporter plasmids containing the entire exonic and intronic DNA sequence spanning *SCN1A* exons 20 to 21 (including 20N) were synthesized as described previously^3^. We generated six plasmids, one with the reference sequence and five with the proband-specific variants.

### Splicing reporter assay

*SCN1A* splicing reporter plasmids (5ug) were transfected in HEK293T cells using Lipofectamine LTX with Plus reagent following the manufacturer’s protocol (Thermo Fisher Scientific #15338100). Cells were harvested after 24 hours using TRIzol Reagent (Invitrogen #15596026) and RNA isolation was performed according to manufacturer’s protocol. RNA concentration and quality was assessed using a NanoPhotometer N60 (Implen). RT-qPCR was used to quantify the amount of 20N inclusion (Exon20:20N and 20N:Exon21 junctions), the canonically spliced form (Exon20:Exon21 junction), and *GAPDH* levels (primers in **Supplementary Table S2**). Reverse transcription and qPCR reactions were prepared using iTaq™ Universal SYBR® Green One-Step Kit on a BIO-RAD CFX Connect™ Real-Time System. We used the ΔΔCt method, with *GAPDH* (first delta) and canonical *SCN1A* transcript (Exon20:Exon21) (second delta). Values were then log transformed to obtain the fold change for 20N inclusion compared to the canonical transcript. Three biological and three technical replicates were performed for each target. Student’s two-tailed *t*-test was used to determine significance (*P ≤ 0.05; **P ≤ 0.01).

### Human iPSC culture

Human induced pluripotent stem cells (iPSCs) were maintained in mTeSR+ medium (Stem Cell Technologies #05825) on tissue culture-treated plates coated with Matrigel (Corning #354277). Two control lines were purchased from Coriell: GM03651 (female) and GM03652 (male).

### Generation of neurons from patient-specific iPSCs (Protocol 1)

iPSCs were differentiated into neural progenitor cells (NPCs) using the STEMdiff SMADi Neural Induction Kit (STEMCELL Technologies; #08581) Monolayer Culture Protocol following the manufacturer’s protocol. NPCs were cultured on Poly-L-ornithine (Sigma; #P4957) and Laminin (Sigma; #L2020)-coated plates and differentiated into neuronal precursors using STEMdiff Forebrain Neuron Differentiation Kit (STEMCELL Technologies; #08600), and the cells were further matured using STEMdiff Forebrain Neuron Maturation Kit (STEMCELL Technologies; #08605). Neurons were maintained in maturation media for 25-35 days, at which time they were harvested in TRIzol reagent (Invitrogen #15596026).

### Generation of neurons from patient-specific iPSCs (Protocol 2)

iPSCs were differentiated into neurons as described previously^22^ with the following modifications: On Day 0, iPSCs were dissociated using Accutase and collected in mTeSR Plus medium (STEMCELL Technologies) containing Y27632 (10 uM, Reprocell) in a microcentrifuge tube. rtTA (5 ul per 1.2×10^6^ cells) and NGN2 (2.5 ul per 1.2×10^6^ cells) lentiviruses were added directly to cell suspension and incubated for 10 minutes at room temperature. Cells were seeded in 10 cm dishes coated with hESC-qualified matrigel (Corning; #354277). On Day 1, the culture medium was replaced with a 1:1 ratio of mTeSR Plus medium and Neurobasal Medium (Gibco) supplemented with Glutamax (Gibco), NEAA (Gibco), N2 (GeminiBio), and Gem21 (GeminiBio). Doxycycline (3 ug/ml, Sigma Aldrich) was added to induce TetO gene expression. On Day 2, culture medium was replaced with Neurobasal Medium containing BDNF (10 ng/ml, PeproTech), NT-3 (10 ng/ml, PeproTech), SB431542 (10 uM, Sigma Aldrich), XAV939 (2 uM, PeproTech), LDN-193189 (100 nM, Tocris), doxycycline (2 ug/ml), and puromycin (1 ug/ml, AlfaAesar) for 72 hours of puromycin selection. Starting from Day 5, cells were maintained in Neurobasal Medium containing BDNF and NT-3 (10 ng/ml each) and doxycycline (2 ug/ml) until harvested for Western blotting and RNA sequencing analysis. In order to eliminate contamination of non-neuronal cells in culture for biochemical assay and RNA sequencing analysis, AraC (2 ug/ml, Sigma Aldrich) was added during media change.

### Membrane preparation and Western blotting

Cultured human iPSC-derived NGN2 neurons were maintained in 6-well plates and were lysed on day 21-24 for membrane protein extraction using the Mem-PER Plus Membrane Protein Extraction kit (Thermo Scientific) according to manufacturer’s instructions. To optimize the yield, permeabilization buffer and solubilization buffer were supplemented with phosphatase inhibitor (Roche) and Halt Protease Inhibitor Cocktail (Thermo Scientific). Solubilized membrane protein samples were quantified using Pierce BCA Protein Assay kit (Thermo Scientific), and the samples were denatured with NuPAGE LDS sample buffer (Invitrogen) and boiling at 70°C for 5 minutes. Protein was separated on 3 to 8% NuPAGE Tris-Acetate gel (Invitrogen) and transferred to PVDF membrane. Membranes were blotted with primary anti-Na_v_1.1 (1:800, Alomone Labs #ASC-001), anti-Na/K-ATPase (1:1000, Addgene #180089) (membrane protein loading control), and anti-Synaptotagmin (1:1000; Proteintech #14511-1-AP) antibodies. Beta-actin (1:1000, Cell Signaling Technologies #3700S) was also used as additional sample loading control. Western blot bands were captured on Odyssey Fc Imaging System (LI-COR Biosciences) and quantified using ImageJ software by normalizing to the membrane protein loading control.

### Targeted long-read RNA-sequencing

Cultured human iPSC-derived neurons (differentiation protocol 1) were harvested in TRIzol reagent (Invitrogen #15596026). Following TRIzol/chloroform extraction, RNA was purified from the aqueous layer using RNA Clean & Concentrator (Zymo #R1017). RNA concentration and quality was assessed by High Sensitivity RNA ScreenTape on a Tapestation 4150 (Agilent). RNA with an RNA integrity number (RIN) greater than 9 was used for library preparation. 1ug RNA was used for cDNA synthesis, which was performed using the Maxima H Minus cDNA Synthesis Master Mix (Thermo Scientific #M1661). 200ng of RNA-equivalent cDNA was amplified with LongAmp Taq 2X Master Mix (NEB #M0287S) and gene-specific primers tailed with Oxford Nanopore universal sequences (**Supplementary Table S3**). 200 femtomoles of PCR 1 product was then amplified with LongAmp Taq 2X Master Mix and barcoding primers (Oxford Nanopore Technologies PCR Barcoding Expansion 1-12 #EXPPBC001). Samples were pooled and the library was prepared for sequencing with the Ligation Sequencing Kit (Oxford Nanopore TechnologiesSQK-LSK109). Libraries were sequenced on an Oxford Nanopore Technologies MinION flow cell.

### Long-read sequencing data analysis

Bases were called from Nanopore data using Guppy. Adapters were trimmed and concatenated reads were separated using Porechop (available at https://github.com/rrwick/Porechop)^23^. Reads were filtered for quality and trimmed using Nanofilt^24^. Reads were mapped to the human (hg38 build) genome using minimap2^25,26^. Transcript analysis was performed using TALON^27^.

### Bulk mRNA sequencing

Cultured human iPSC-derived neurons (differentiation protocol 1) were harvested and RNA was prepared and quantified as described above. RNA with an RIN greater than 8 was used for library preparation. mRNA enrichment libraries were prepared and sequenced with paired-end 150 bp reads on an Illumina platform at a depth of ∼30 million reads per sample.

### Bulk mRNA sequencing analysis

Adapters were trimmed with HTStream (available at https://github.com/s4hts/HTStream). Sequencing reads were mapped to the human (hg38 build) genome using STAR^28^ and data were normalized in DESeq2^29^.

### RNA antisense purification coupled with mass spectrometry (RAP-MS)

RAP–MS was carried out as described previously (McHugh et al. Nature 2015) with the following modifications: RNA antisense probes were designed using the RAPOligoDesigner software to capture mature and immature mRNA transcripts (**Supplementary Table S4**) and probes were synthesized with a 5’ biotinylated modification. HEK293T cells were transfected with pDESTSplice reporter plasmids using Lipofectamine LTX with Plus reagent (Thermo Fisher Scientific #15338100). Two replicates were prepared for each plasmid. Twenty-four hours post transfection, media was replaced with cold phosphate-buffered saline (PBS) and RNA-RBP interactions were covalently captured by UV-crossinkling using a UV Stratalinker 1800 (Stratagene), followed by nuclear fractionation, RNA antisense purification, and protein harvesting. As a negative control, the same procedure was performed in untransfected cells. Proteomics analysis was performed using Orbitrap Exploris 480.

Proteomics data was analyzed in the Scaffold version 5 software and proteins identified against the human NCBI proteome database (Uniprot-SP-human database (20190906_20200629)) with a protein threshold of 99% and a minimum of 2 peptides at a threshold of 95%. To remove false positives, we excluded all proteins that were also recovered in the GFP-only plasmid transfected cells or untransfected cells.

### RNA Immunoprecipitation (RIP)

*SCN1A* splicing reporter plasmids (5ug) were transfected in HEK293T cells using TurboFectin 8.0 (Origene #TF81001). After twenty-four hours, UV-crosslinking was performed as above and cell pellets were flash-frozen using liquid nitrogen. Cell pellets thawed on ice and lysed with RIPA Lysis and Extraction Buffer (Thermo Scientific #89901) prepared with cOmplete EDTA-free Protease Inhibitor Cocktail (Roche #11836170001). Lysates were centrifuged at 14,000 x g for 15 minutes at 4°C, and the supernatant was used for immunoprecipitation. Dynabeads Protein A or Protein G magnetic beads (Thermo Fisher Scientific #10001D, #10003D) were prepared and immunoprecipitation was performed using 1-5ug of antibody and a ten- or thirty-minute IP incubation (antibodies and details in **Supplementary Table S5**). After washing, RNA was eluted with a nondenaturing elution and treated with 2ug Proteinase K (Qiagen #19131) for 15 minutes at 55°C followed by the addition of TRIzol Reagent (Invitrogen #15596026) and isolated as above. RNA concentration and quality was assessed by High Sensitivity RNA ScreenTape on a Tapestation 4150 (Agilent). One representative sample was used to assess antibody specificity. RIP was carried out with antibodies for RBPs of interest or an NMS control. Following the immunoprecipitation incubation, the supernatant was saved for Western blot analysis. After washing, each sample was separated for protein and RNA analysis (**Supplementary Figure S3**). Supernatant and eluted protein samples were prepared for Western blot with a denaturing NuPAGE LDS Sample buffer (Invitrogen #NP0007).

### ddPCR with immunoprecipitated RNA

cDNA was synthesized from 100ng immunoprecipitated RNA using the iScript Reverse Transcription Supermix (Bio-Rad) according to the manufacturer’s instructions. 20ng of RNA-equivalent cDNA was used as template for droplet digital PCR (ddPCR) which was performed as described above. Each sample was assayed for *SCN1A* 20N inclusion (20N:Exon21 junction) and the canonically spliced form (Exon20:Exon21 junction) (primers in **Supplementary Table S6**), with 3-4 biological replicates per group. Kruskal-Wallis comparisons with Dunn’s post-hoc correction were used to determine significance (*P ≤ 0.05; **P ≤ 0.01).

## Ethics declarations

I.E.S. has served on scientific advisory boards for BioMarin, Chiesi, Eisai, Encoded Therapeutics, GlaxoSmithKline, Knopp Biosciences, Nutricia, Rogcon, Takeda Pharmaceuticals, UCB, Xenon Pharmaceuticals; has received speaker honoraria from GlaxoSmithKline, UCB, BioMarin, Biocodex, Chiesi, Liva Nova, Nutricia, Zuellig Pharma and Eisai; has received funding for travel from UCB, Biocodex, GlaxoSmithKline, Biomarin, Encoded Therapeutics and Eisai; has served as an investigator for Anavex Life Sciences, Cerecin Inc, Cerevel Therapeutics, Eisai, Encoded Therapeutics, EpiMinder Inc, Epygenyx, ES-Therapeutics, GW Pharma, Marinus, Neurocrine BioSciences, Ovid Therapeutics, Takeda Pharmaceuticals, UCB, Ultragenyx, Xenon Pharmaceuticals, Zogenix and Zynerba; and has consulted for Care Beyond Diagnosis, Epilepsy Consortium, Atheneum Partners, Ovid Therapeutics, UCB, Zynerba Pharmaceuticals, BioMarin, Encoded Therapeutics and Biohaven Pharmaceuticals; and is a Non-Executive Director of Bellberry Ltd and a Director of the Australian Academy of Health and Medical Sciences and the Australian Council of Learned Academies Limited. She may accrue future revenue on pending patent WO61/010176 (filed: 2008): Therapeutic Compound; has a patent for *SCN1A* testing held by Bionomics Inc and licensed to various diagnostic companies; has a patent molecular diagnostic/theranostic target for benign familial infantile epilepsy (BFIE) [PRRT2] 2011904493 & 2012900190 and PCT/AU2012/001321 (TECH ID:2012-009). The remaining authors have nothing to disclose.

## Acknowledgements

We thank the patients and their families for participating in our research study. We also thank Jennifer Kearney for her sodium channel expertise, which was invaluable for integrating the long-read and short-read sequencing analyses. Thank you to Jennifer Ching Man Wai for her valuable assistance with long read sequencing. Long Read Sequencing was supported, in part, by the Northwestern University NUSeq Core Facility. The Northwestern Proteomics Core Facility is supported by NCI CCSG P30 CA060553 awarded to the Robert H Lurie Comprehensive Cancer Center, instrumentation award (S10OD025194) from NIH Office of the Director and the National Resource for Translational and Developmental Proteomics supported by P41 GM108569. We also thank Kwang-Youn Kim for lending his expertise in genetic statistics. This study was supported by the Dravet Syndrome Foundation Research Grant and NIH NINDS R21NS121572 both to G.L.C.

## Contributions

Conceptualization: H.C.H., P.N.S., H.C.M., G.L.C.; Methodology: H.C.H., P.N.S., J.H., J.D.C.; Software: C.G.B., F.R.; Resources: K.E., S.W., I.E.S., Validation: H.C.H.; Formal Analysis: H.C.H., J.H., C.G.B., J.D.C., G.L.C.; Investigation: H.C.H., P.N.S., J.H., E.G., M.B., A.R.; Visualization: H.C.H.; Writing – Original draft: H.C.H., G.L.C.; Writing – Review & Editing: All authors; Supervision: G.L.C.; Funding acquisition: G.L.C.

