## Supplementary Information for "Long-read sequencing and profiling of RNA-binding proteins reveals the pathogenic mechanism of aberrant splicing of an *SCN1A* poison exon in epilepsy"

**Figure S1. Exon 20N inclusion measured by splicing reporter assay in HEK293T cells.** WT or proband-specific splicing reporter plasmids containing either WT sequence or proband-specific variant were transfected into HEK293T cells in triplicate and harvested after 24 hours for RNA. RT-qPCR was used to measure the amount of canonical *SCN1A* (Exon20:Exon21), 20N inclusion (20N:Exon21 and Exon20:20N). *GAPDH* was measured as a housekeeping gene. Cycle threshold (Ct) values were normalized to *GAPDH* ($\Delta$Ct) and then 20N:Exon21 or Exon20:20N was normalized to Exon20:Exon21 ($\Delta\Delta$Ct). $\Delta\Delta$Ct values were log transformed to obtain the fold change of 20N inclusion compared to canonical transcript. Compared to WT, variants 2 and 3 lead to increased 20N inclusion as measured by both assays, and variant 1 leads to increased 20N inclusion as measured by the 20N:Exon21 assay. Biological replicates n=3, technical RT-qPCR replicates n=3. qPCR data are presented as the mean ± SD. For comparisons, student's unpaired two-tailed t-test was performed (*P ≤ 0.05; **P ≤ 0.01).

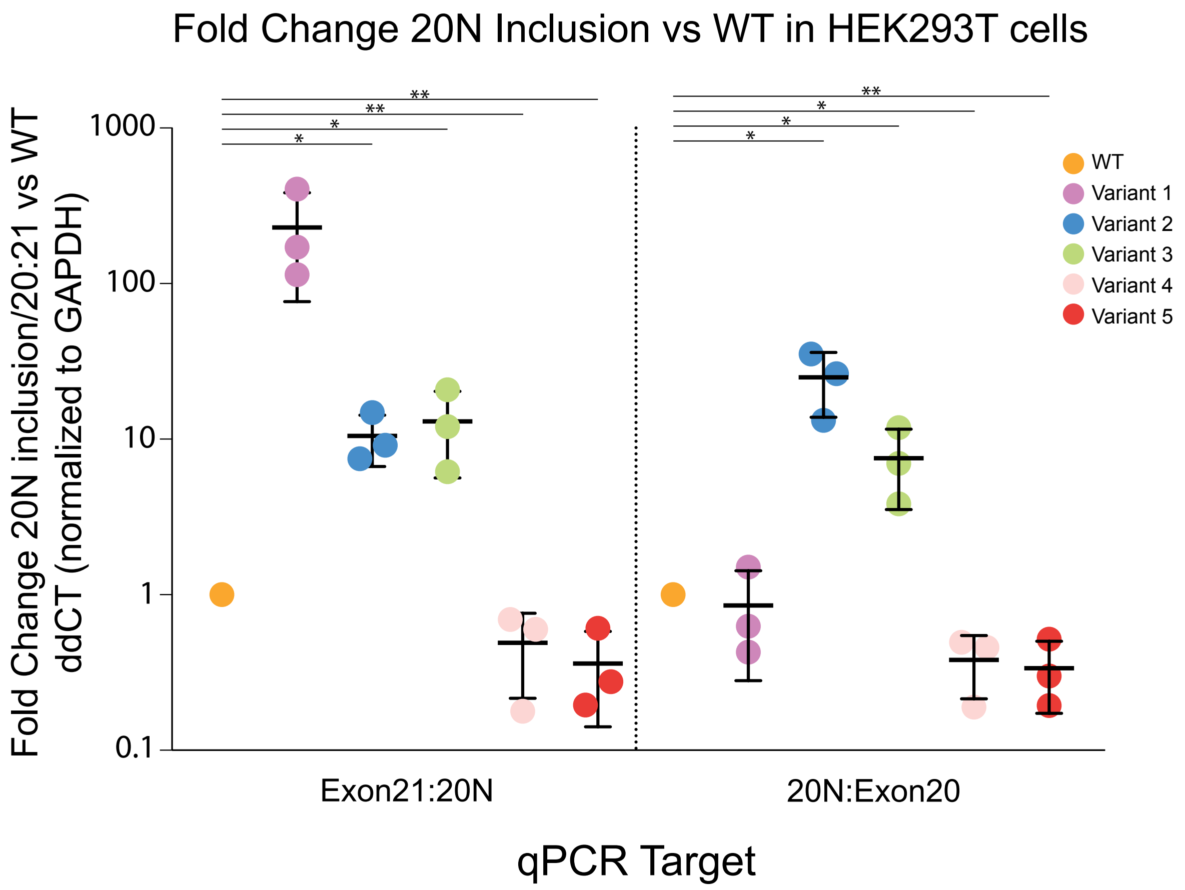

**Figure S2. 20N inclusion introduces a PTC. (A)** A schematic representation of the *SCN1A* exons 20 and 21 in the mature mRNA transcript. **(B)** When *SCN1A* 20N (purple) is included in the mature mRNA transcript, a PTC (red) is introduced in exon 21. **(C)** The mRNA sequence of exon 20, 20N, and exon 21 are shown in codon blocks. When 20N is spliced between exons 20 and 21, the open reading frame is shifted, introducing a PTC in exon 21 (red). This PTC is not located in the final or penultimate exon, therefore this transcript will likely be targeted for NMD.

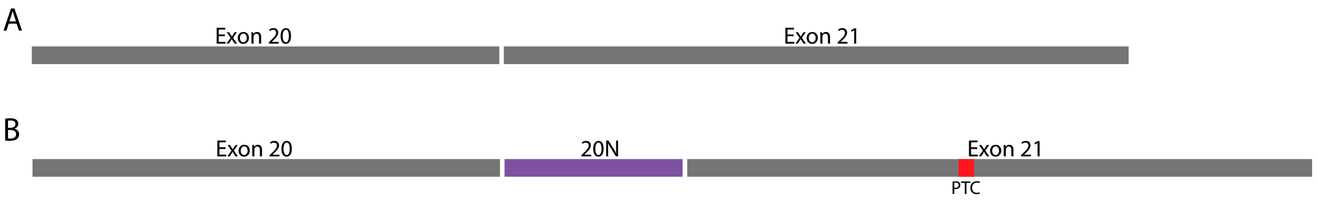


C

GUU UCA UUG GUC AGU UUA ACA GCA AAU GCC UUG GGU UAC UCA GAA CUU GGA GCC AUC AAA UCU CUC AGG
ACA CUA AGA GCU CUG AGA CCU CUA AGA GCC UUA UCU CGA UUU GAA GGG AUG AGG GAU AAU CUU GCU CCA
ACU UGG AUG GGG UGG AGC GCU GGU UCC UCC CCU GAG CCC UUU AUU AUG GGU GGU UGU GAA UGC CCU UUU
AGG AGC AAU UCC AUC CAU CAU GAA UGU GCU UCU GGU UUG UCU UAU AUU CUG GCU AAU UUU CAG CAU CAU
GGG CGU AAA UUU GUU UGC UGG CAA AUU CUA CCA CUG UAU UAA CAC CAC AAC UGG UGA CAG GUU UGA CAU
CGA AGA CGU GAA UAA UCA UAC UGA UUG CCU AAA ACU AAU AGA AAG AAA UGA GAC UGC UCG AUG GAA AAA
UGU GAA AGU AAA CUU UGA UAA UGU AGG AUU UGG GUA UCU CUC UUU GCU UCA AGU U

**Figure S3. Antibodies used for RNA-immunoprecipitation specifically pull-down protein of interest and RNA.** Splicing reporter plasmids were transfected into HEK293T cells. Twenty-four hours after transfection, RNA-protein interactions were covalently captured by UV-crosslinking. RIP was carried out with antibodies for RBPs of interest or an NMS control. Following the immunoprecipitation incubation, the supernatant was saved for Western blot analysis (Supernatant). After washing, each sample was separated for protein (A, C) and RNA analysis by Tapestation (B, D). One representative sample was used to assess RIP specificity. **(A)** Anti-SRSF1 antibody specifically pulls down SRSF1, as indicated by the 28kD and 33kD bands in samples incubated with antibody and eluted from the IP. Samples incubated with NMS instead of antibody results in no pulldown of SRSF1 in the eluted sample and presence in the IP supernatant. **(B)** RNA is only detected in samples incubated with anti-SRSF1 antibody. **(C)** Anti-HNRNPL antibody specifically pulls down HNRNPL, as indicated by the 60kD and 66kD bands in samples incubated with antibody and eluted from the IP. Samples incubated with NMS instead of antibody results in the absence of HNRNPL in the eluted sample and presence in the IP supernatant. **(D)** RNA is only detected in samples incubated with anti-HNRNPL antibody. HC: heavy chain; NMS: normal mouse serum; nt: nucleotide; kD: kilodalton.

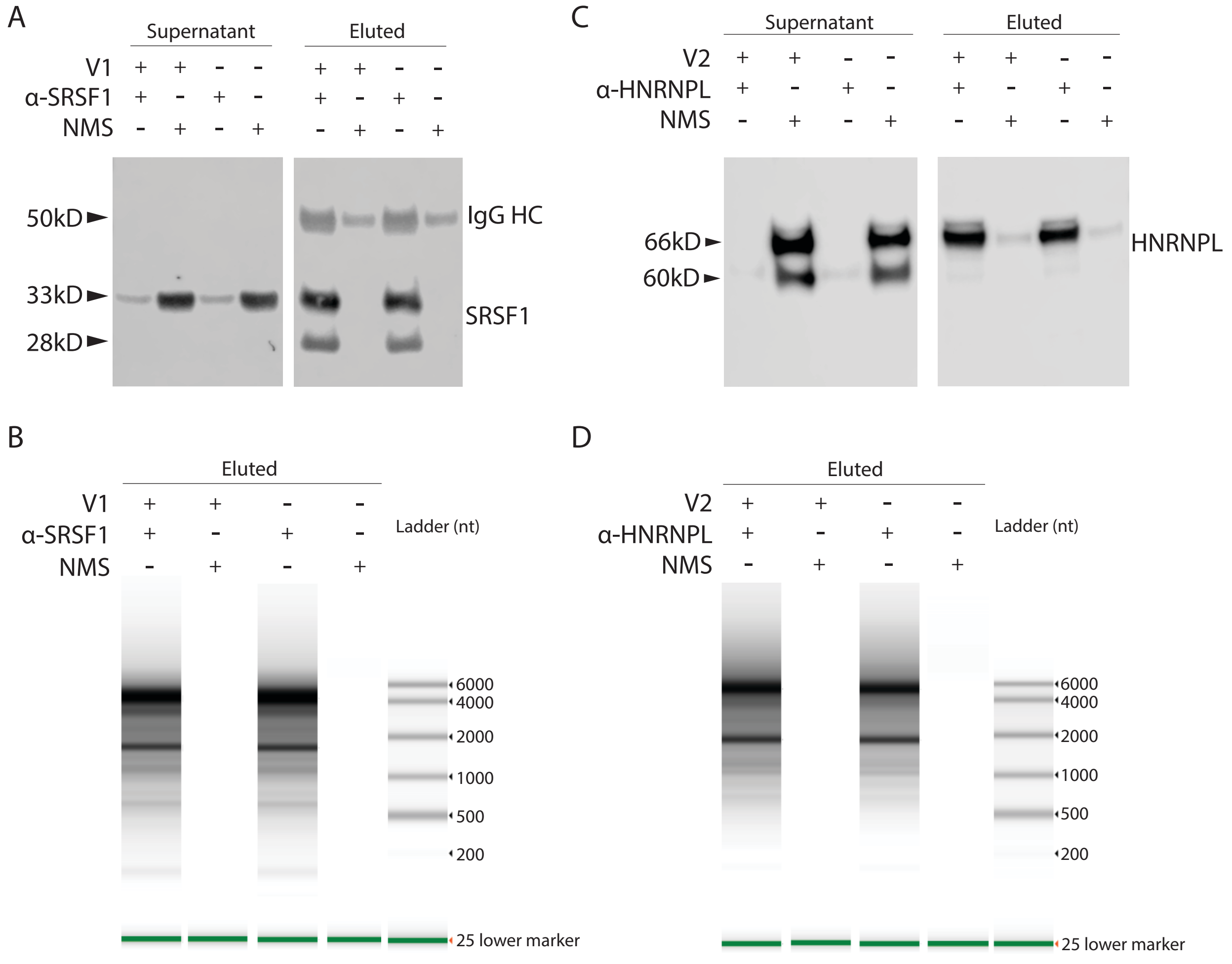

**Figure S4. RIP and ddPCR for canonical *SCN1A* transcript.**

**(A and B)** RIP was performed with the same splicing reporter constructs used for RAP-MS with the addition of a variant 3 construct. RNA-RBP interactions were captured by UV cross-linking, and immunoprecipitations were carried out with antibodies for SRSF1 (A) or HNRNPL (B). Following RNA extraction and reverse transcription, droplet-digital PCR with 20ng RNA was used to quantify canonical transcripts bound to SRSF1 (A) or HNRNPL (B). Kruskal-Wallis comparisons with Dunn's post-hoc test were performed on WT-variant pairs (SRSF1) or variant 1-WT/variant pairs (HNRNPL) (*P-adjusted ≤ 0.05; **P ≤ 0.01). Biological replicates *n* = 4 or *n* =2 (HNRNPL variant 3), technical replicates *n* = 2. Data are presented as the mean ± SD.

A

B

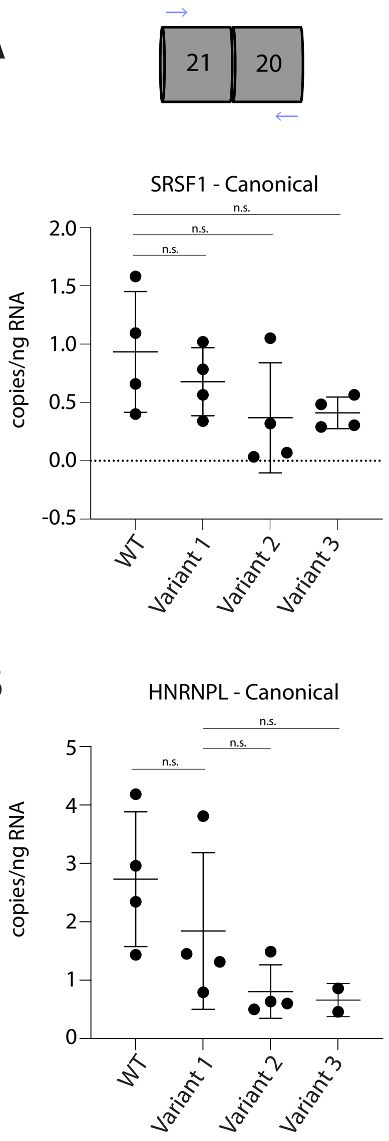

**Figure S5. Variants alter RBP motifs in mRNA.** Putative motifs are highlighted in gray. **(A)** Variant 1 changes a C to a U, disrupting a CA motif. **(B)** Variants 2 and 3 both disrupt G residues. Variant 3 causes a G-to-C (mRNA) change, which may enhance SRSF1 binding. The mechanism by which variant 2 enhances SRSF1 binding is less clear. **(C)** Variants 4 and 5 create AU-rich motifs (AAAUUUU), which are recognized by ELAVL3 and ELAVL4.

A. WT 5′–AUUACCCCUUUUG–3′
HNRNPL
V1 5′–AUUACCUCUUUUG–3′

B. WT 5′–GGAUGGGGUGGAGCGCT–3′
V2 5′–GGAUGGCGUGAAGCGCT–3
SRSF1
V3 5′–GGAUGGCGUGAAGCGCT–3
SRSF1

C. WT 5′–AAAUUUCCAGUCCAAAUUUU–3′
V4 5′–AAAUUUCA––––––AAUUUU–3′
ELAVL3/4
V5 5′–AAAUUUUCAGUCCAAAUUUU–3′
ELAVL3/4

**Table S1. Variant details and ACMG criteria/classification.**

| **Proband** | **Variant description – Genomic** NC_000002.12 | **Variant description – Transcript** NM_001165963.4: | **ACMG criteria** | **ACMG classification** |
|---|---|---|---|---|
| **1** | g.166007216G>A | c.4002+2503C>T | PVS1, PS4_mod, PM2, PP1, PP3 | Pathogenic |
| **2** | g.166007264C>T | c.4002+2455G>A | PVS1, PS2, PS4_mod, PM2, PP3 | Pathogenic |
| **3** | g.166007268C>G | c.4002+2451G>C | PVS1, PS2, PS4_mod, PM2, PP3 | Pathogenic |
| **4** | g.166007551_166007555del | c.4002+2168_4002+2172del | PVS1, PS4_mod, PM2, PP3 | Pathogenic |
| **5** | g.166007554G>A | c.4002+2165C>T | PVS1, PS4_mod, PM2, PP1, PP3 | Pathogenic |

*We previously reported[3] all five probands/variants in the opposite numeric order

**Table S2. Splicing assay qPCR primer details.**

| **Primer name** | **Forward sequence (5'-3')** | **Reverse sequence (5'-3')** |
|---|---|---|
| GAPDH_Exon_3:Exon_4 | TCAATGGAAATCCCATCACC | TTGATTTTGGAGGGATCTCG |
| SCN1A_Exon_20:Exon_21 | GGATGAGGGTGGTTGTGAAT | TCATGATGGATGGAATTGCT |
| SCN1A_Exon_20N:Exon_21 | CCTCCCCTGAGCCCTTTAT | AAAAGGGCATTCACAACCAC |
| SCN1A_Exon_20:Exon_20N | GAGACCTCTAAGAGCCTTATCTCG | GCTCAGGGGAGGAACCAG |

**Table S3. Long read sequencing primer details.**

| **Primer name** | **Forward sequence (5’-3’)** | **Reverse sequence (5’-3’)** |
|---|---|---|
| Scn1a_h_202 | TTTCTGTTGGTGCTGATATTGCGGCCCAAAGCCAAATAGTGAC | ACTTGCCTGTCGCTCTATCTTCCAACGCAGGAAGGGACATCA |

**Table S4. RAP-MS probes.**

| Probe number | Sequence (5’-3’) |
|---|---|
| 1 | CCTTGCACTGTGACTTATGTGTAGTGGGGTGAGGGAGGGATTGGGAAGGGTACTATTATTGCACCACAGTAGGGAAAATACATTATTTAC |
| 2 | TTTTGTATAGGATAATCTTGCTCCAACTTGGATGGGGTGGAGCGCTGGTTCCTCCCCTGAGCCCTTTATTATGGGTACTGTATTACCCCT |
| 3 | AACTATATATTTCTATTTTGTATAGGATAATCTTGCTCCAACTTGGATGGGGTGGAGCGCTGGTTCCTCCCCTGAGCCCTTTATTATGGG |
| 4 | AGTCTTAATGGATAGGGAAACAGACGGAGAGCATCATGAACAAAAAGTAACACCAAATGTTCTGTCATATCAGATTTCTAACTAATAACA |

**Table S5. RNA immunoprecipitation antibody details.**

| **Antibody** | **Vendor (Catalog number)** | **Protein A/G** | **Amount used for immunoprecipitation (ug)** | **IP duration (minutes)** | **Dilution used for Western Blot** |
|---|---|---|---|---|---|
| ASF/SF2 monoclonal antibody | Proteintech (#66671-1-Ig) | G | 5 | 10 | 1:1000 |
| hnRNP L (E5J7B) Rabbit monoclonal antibody | Cell Signaling Technologies (#65043) | A | 1 | 30 | 1:1000 |

**Table S6. ddPCR primer details.**

| Target amplicon | Forward sequence (5'-3') | Reverse sequence (5'-3') |
|---|---|---|
| SCN1A_Exon_20:21 | AGACAAACCAGAAGCACATTCA | GAGACCTCTAAGAGCCTTATCTCG |
| SCN1A_Exon_20N:21 | AGACAAACCAGAAGCACATTCA | CTGGTTCCTCCCCTGAGC |
